# Modeling cellular influence delineates functionally relevant cellular neighborhoods in primary and metastatic pancreatic ductal adenocarcinoma

**DOI:** 10.1101/2025.06.12.659314

**Authors:** Yeonju Cho, Jae W. Lee, Sarah M. Shin, Alexei G. Hernandez, Xuan Yuan, Jowaly Schneider, Jody E. Hooper, Laura D. Wood, Elizabeth M. Jaffee, Atul Deshpande, Won Jin Ho

## Abstract

Pancreatic ductal adenocarcinoma (PDAC) is a highly lethal cancer, with liver metastases significantly worsening outcomes. However, distinct features of the tumor microenvironment (TME) between primary and metastatic sites remain poorly defined. Cellular neighborhoods within the TME are recognized as functional units that influence tumor behavior. Conventional spatial methods, which assign equal weights to all cells in a region, fail to capture the nuances of cellular interactions. To address this, we developed Functional Cellular Neighborhood (FunCN) quantification, which integrates both the proportion and proximity of surrounding cells. Applying FunCN to PDAC imaging mass cytometry data, we identified neutrophil-enriched interactions in liver metastases compared to primary tumors, correlating with elevated VISTA expression by tumor cells. Additionally, FunCN clusters around CD8^+^ T cells in pancreas and liver were associated with higher TIGIT and LAG3, respectively. These findings demonstrate the importance of spatial immune landscapes in PDAC and identify potential therapeutic opportunities.

## Introduction

Pancreatic ductal adenocarcinoma (PDAC) is among the most deadliest malignancies^1^, largely due to its propensity to metastasize early to distant organs and poor response to existing treatments^2^. More than 50% of PDAC patients present with liver metastases at diagnosis, which are associated with particularly poor prognosis.^3^ Thus, understanding mechanisms that enable tumor cells to spread and proliferate at distant metastatic sites, including the liver, is crucial for identification of novel therapeutic targets that may improve patient outcomes.

Recent studies have shown that primary and metastatic tumors are heterogeneous and that the tumor microenvironment (TME) plays a key role in modulating cancer progression and metastasis. The TME is defined not only by its cellular components but also by interactions between tumor cells and non-tumor cells, including immune cells and fibroblasts, that can support or suppress tumor growth^4^. In fact, the distance between immune and tumor cells directly reflects both the effectiveness of immune cells in eliminating tumor cells and the tumor’s capacity to influence immune cells.^5^ For example, a recent study^6^ demonstrated that tumors with PD-1^+^ T cells located within approximately 10 µm of PD-L1^+^ tumor cells exhibited significantly longer progression-free survival following PD-1 blockade therapy, highlighting the importance of assessing spatial interactions when evaluating therapeutic outcomes. Previous studies^7–9^ have integrated cellular proximity and influence by taking into consideration distances between cells and the cell density within specific regions. However, these methods impose rigid spatial thresholds – either by fixing the number of nearest neighbors or by categorizing interactions using predefined ranges (e.g., 10-20 µm, 20-30 µm) – which may overlook biologically relevant cells just beyond these cutoffs. Moreover, they assign equal weights to all included cells, regardless of their precise distance. This uniform weighting limits the resolution and accuracy of spatial interaction analysis within the TME.

To address these limitations and achieve a more comprehensive characterization of the TME, we developed a novel method called Functional Cellular Neighborhoods (FunCN) that quantifies cellular interactions by considering both the proximity and frequency of neighboring cells. Specifically, FunCN uses a spatial kernel-based approach to model the decrease in a cell’s influence on other cells as they get farther apart. We applied this approach to investigate the TME of primary and metastatic PDAC and identified distinct neighborhoods of CD8^+^ T cells and tumor cells. We also found cellular interactions that were unique to the pancreas and liver. Overall, our study provides insight into the cellular interactions within primary and metastatic tumors that may guide future therapeutic strategies.

## Method

### Human samples

Primary and metastatic tumors were obtained from patients with metastatic PDAC through the Johns Hopkins Research Autopsy Program. Written informed consent was obtained from all research subjects, and the research protocol was approved by the institutional review board of the Johns Hopkins School of Medicine (protocol number NA_00036610). Tissue collection was carried out in compliance with the 1996 Declaration of Helsinki.

Collected tissue specimens were fixed in formalin and embedded in paraffin. Tissue sections were then stained with hematoxylin and eosin and reviewed by board certified anatomic pathologists (J.E.H. and L.D.W.). Regions containing tumor cells were then selected to construct tissue microarrays (TMAs) with each core measuring approximately 1 mm in diameter. Patient characteristics and the number of cores analyzed from each patient are shown in Supplementary Table 1.

### IMC staining and IMC processing

IMC staining on TMAs was performed as previously described^10^. Briefly, the TMA slides were baked, deparaffinized in xylene, and rehydrated in an alcohol gradient. Heat-mediated antigen retrieval was performed at 95 °C for 30 minutes using Antigen Retrieval Agent pH9 (Agilent S2367). The slides were then blocked with 3% BSA for 45 min at room temperature, followed by overnight staining at 4 °C with the antibody cocktail. Information on antibodies used for IMC is shown on Supplementary Table 2. After staining, images were acquired using the Hyperion Imaging System (Standard BioTools) at the Johns Hopkins Mass Cytometry Facility. Images were visualized using MCD Viewer (Standard BioTools). For data analysis, image segmentation was carried out using CellProfiler^11^ (version 3.1.9), ilastik^12^ (1.4.1rc2-gpu), and HistoCAT^13^ (version 1.76).

### Data pre-processing

The data was pre-processed with the removal of benign cells, followed by log-normalization of the expression data (log2(expr + 1)). We performed batch effect correction using the Harmony^14^ R package (version 1.2.3), accounting for patient and tissue site differences. The adjusted expression values were subsequently shifted to ensure non-negative values by subtracting the minimum expression level for each marker.

### Cell type annotation

#### Cell clustering

We performed clustering using a Self-Organizing Map (SOM), which organized the high-dimensional data into a 35×35 grid, resulting in 1,225 nodes with using FlowSOM^15^ R package (version 2.12.0). Cell-type specific markers (CD3, CD8, FOXP3, CD4, granzyme B (GZMB), CD45RA, CD57, DCSIGN, CD15, CD86, CD163, CD206, CD68, HLA-DR, CD74, VISTA, COL1A1, smooth muscle actin and vimentin (SMA/VIM), pan-keratin (CK), and Ki-67) from the panel (**Figure S1A**) were selected for clustering. The cell expression values within each node were summarized by their median expression of the cells in that node, and outliers were controlled by applying a median absolute deviation (MAD) threshold of 2 median absolute deviations (MADs). Subsequent metaclustering of the SOM nodes using Consensus Clustering was performed with ConsensusClusterPlus R package (version 1.68) to identify distinct 50 clusters using Euclidean distance

#### Visualization and annotation

To facilitate robust cell type annotation, we generated a heatmap visualization of the protein expression for each cluster. The expression data was normalized using min-max scaling to a [0, 1] range, based on the 33rd and 99th percentiles for heatmap visualization.

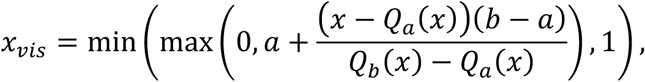

where *Q_a_*(*x*) and *Q_b_*(*x*) are the *a*^th^ and *b*^th^ quantile values of the expression data *x* before normalization. We used *a* = 0.33, *b* = 0.99 to highlight positive outliers of protein expressions compared to a symmetric mapping and consequently identify distinct cell types. This visualization improvement is especially beneficial in cases where data sparsity and a skewed distribution could lead to spurious normalization effects. For instance, setting *a* = 0.01, *b* = 0.99 for a highly skewed protein expression distribution results in lower quantiles (*x* < 0.5) of expression values to map to (*x_vis_* > 0.5), resulting in a distorted and misleading visualizations of the expression values. Since we primarily use overexpression of proteins to identify cellular phenotypes, the more robust mapping from *x* → x_vis_ for *x* < 0.5 obtained using *a* = 0.33, allowing for a more accurate cell type annotation. The normalized values were summarized within each cluster, and hierarchical clustering was applied.

#### Subtype annotation and statistical analysis of cellular abundance

After annotation, subtypes of CD4^+^ T cells, other myeloid cells (excluding neutrophil cell type) and stromal cells were identified using the same clustering method but with a 10×10 grid. The heatmap was then generated using ComplexHeatmap^16^ R package (version 2.20) in R to visualize the clustering result. The abundance of each cell type was visualized using bar plots generated using ggplot2 package (version 3.5.1) in R for each tissue site, core, and patient. To evaluate differences in cell-type densities across tissue sites, an unpaired Wilcoxon rank-sum test was performed using the ggpubr package (version 0.6) in R. The resulting median values and standard deviations were then illustrated on scatter plots.

### Functional cellular neighborhood (FunCN) quantification

We present our novel method for quantifying functionally relevant cellular neighborhoods as observed by each individual cell in the spatial dataset. Previous approaches define a cell’s neighborhood either by including all cells within a given radius *r* from a reference cell (distance based CN^17,18^), or by selecting the k-nearest neighbors of the reference cell (knn-based CN^19^), and summarize the proportion of each cell type among the neighboring cells. However, these methods have limitations. First, they do not account for the relative distance of neighboring cells from the reference cell, treating nearby and farther cells equally within the boundary. Second, both methods place a definite boundary on the cellular neighborhood and hence do not allow us to model the influence of cells situated beyond a radius r or beyond the k-th nearest neighbor, respectively. For example, a large tertiary lymphoid structure (TLS) at a moderate distance from a cell may have a non-trivial influence on the cell’s function, despite being outside the boundaries described in the prior art. To overcome these limitations, our novel method for Functional Cellular Neighborhood (FunCN) quantification uses a spatial kernel-based approach to model the influence of all other surrounding cells weighted by their proximity to the reference cell. To quantify the FunCN of the *i*-th cell *C_i_*, we calculate a kernel-weighted influence from its surrounding cells

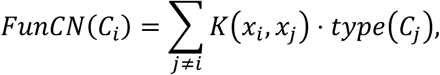

where *type*(*C_j_*) is the cell-type of the *j*-th cell *C_j_* (*j* ≠ *i*). *x_i_* and *x_j_* are the coordinates of the *i*-th and *j*-th cells, and *K*(*x_i_*, *x_j_*) is a spatial kernel function which decreases as the distance between *x_i_* and *x_j_* increases. In this paper, we define *K* as a 2-D Gaussian kernel

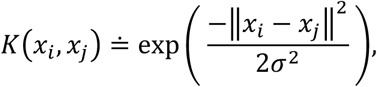

where ‖·‖^+^ is the Euclidean distance, and *σ* = 10*μm* is the width of the kernel. The bandwidth was chosen to place greater emphasis on nearby cells, a critical distance range associated with improved patient survival due to enhanced T cell infiltration within a 10 μm range of tumor^6,20^. Since *K*(*x_i_*, *x_j_*) decreases with increasing intercellular distance between the *i*-th and *j*-th cell, the *j*-th cell will have a smaller influence in the FunCN quantification described above. The resulting FunCN of each cell is thus a weight vector of length equal to the number of distinct cell types *N_celltype_*, with nonnegative weights summing to 1. We used the spatstat^21^ R package (version 3.3.1) to evaluate the weights and subsequently the weighted influence of each cell type for every cell in the dataset (see github repository https://github.com/DeshpandeLab/Spatial_Influence).

### Comparison of FunCN and KNN based cellular neighborhoods

FunCN was compared to the conventional k-nearest neighbor (KNN) approach to assess differences in how cellular proximity is incorporated into influence quantification. To compute the KNN for cellular influences, the FNN R package (version 1.1.4.1) was used to identify 10-nearest neighbors for each cell based on spatial coordinates, with the proportion of the distinct cell types among the 10 nearest neighbors to the reference cell representing its CN. Spatial density of cell counts was visualized using the spatstat^22^ R package. Comparison of FunCN and KNN method were visualized using dot plots, violin plots and histograms using ggplot2.

### Cellular Neighborhood Clustering and Downstream Analysis

Cells were grouped by clustering their FunCN vectors using self-organizing map (SOM) clustering, which grouped cells with similar local microenvironment. To characterize the microenvironment of each CN cluster, we visualized the clusters using dot plot where dot size represented the median cellular influences of each cell type within a cluster and dot color indicated the scaled median value across CNs. Functional marker expression was further evaluated within each clustered CN using radial plots which displayed the scaled median expression of each marker across CNs, relating variations in local microenvironments to functional cell states. To examine trends in functional marker expression across CN clusters, line plots were generated showing absolute median expression values with 95% confidence intervals (calculated as median ± 2 × standard error) for each site. Core 43, which contained approximately 50% immune cell mixture, was excluded from the downstream analysis.

### Survival analysis

Transcriptome and clinical data were obtained from the Cancer Genome Atlas (TCGA) PAAD dataset using TCGAbiolinks package (version 2.32) in R. Primary tumor samples were selected, and raw count data were normalized using the variance stabilizing transformation (VST) from DESeq2 package (version 1.44). Genes with low expression (sum total less than ten reads across all samples) were filtered out. The transformed expression values were used for further statistical analysis including stratification based on gene expression levels, where patients were stratified into high and low expression group based on the 20^th^ percentile threshold. Survival analysis was performed using the survival (version 3.8.3) and survminer (version 0.5) packages to fit and visualize Kaplan-Meier survival curve, assessing survival difference between the high and low gene expression groups.

## Results

### Influence-weighted cellular neighborhoods incorporate proximity and frequency of neighboring cells to enhance characterization of the TME

We present Functional Cellular Neighborhood (FunCN), a novel method for quantifying the cellular neighborhoods using a spatial kernel-based method to model cellular influence in the TME. In our approach, we applied a 2D spatial kernel to model how cellular influence on surrounding cells decays with increasing distance. Spatial kernels (mathematical functions) have been previously used to model influence of cells or biological patterns in spatially resolved transcriptomics^23^. Extending beyond regional-scale analyses, we built on this concept to characterize FunCN for each individual cell, represented as a vector capturing the accumulated influence by all cell types within the tissue.

Applying this framework to map the cellular architecture of PDAC across different sites, we utilized imaging mass cytometry (IMC) with a 35-antibody panel to generate high-dimensional spatial data. A total of 267,239 cells from 64 tissue cores were analyzed, collected from 12 unique patients. This included liver samples from 10 patients and pancreatic samples from 8 patients, with matched primary and metastatic tumors collected from 6 patients. After image processing and cell segmentation, cell types were annotated by clustering cells based on lineage marker expression (**Figure 1A**).

**Figure 1.**
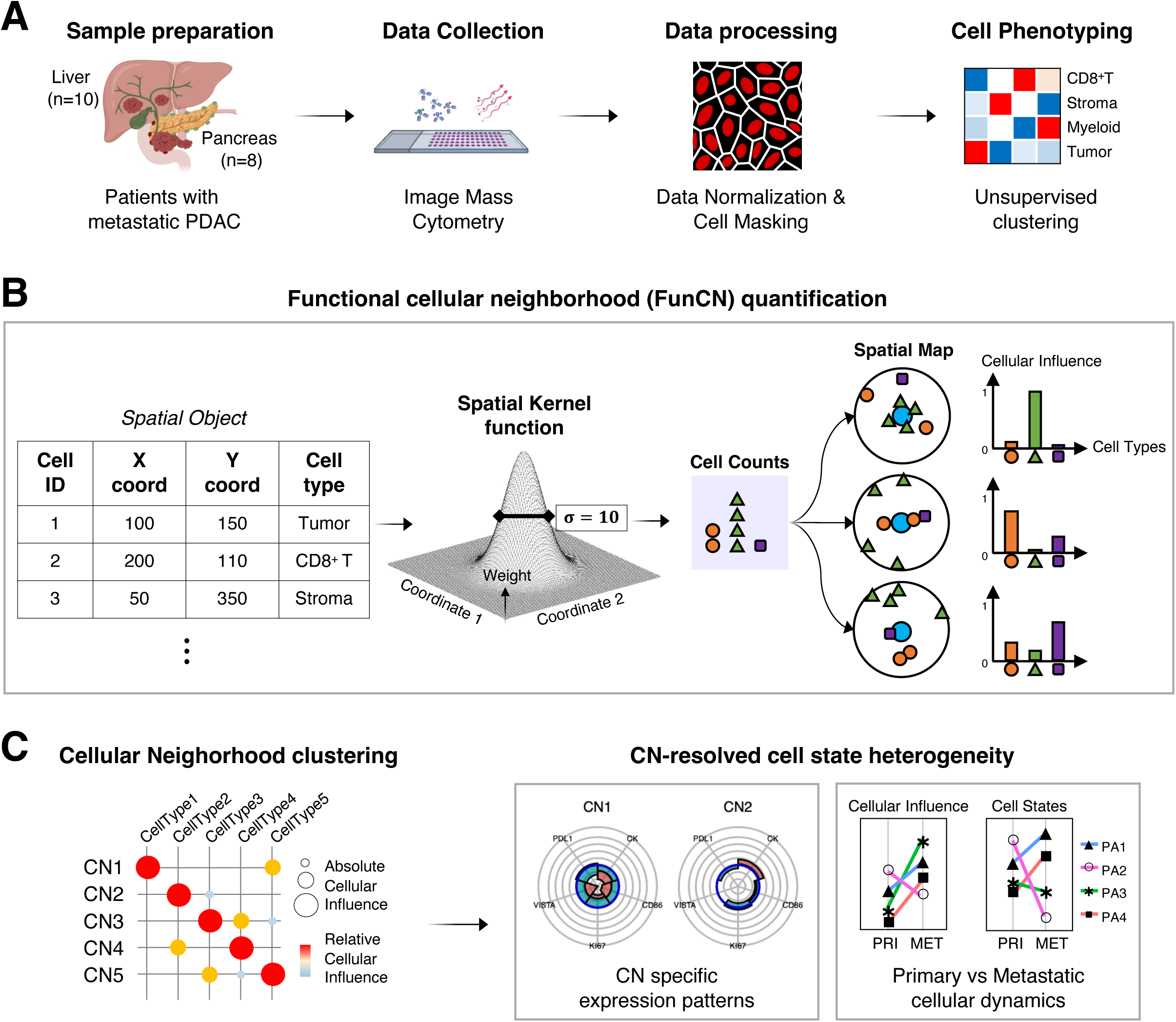
Overview of analysis workflow. **(A)** Metastatic PDAC samples from the liver (*n* = 10) and pancreas (*n* = 8) spatial data were acquired using imaging mass cytometry, followed by data processing and cell type phenotyping. The schematic shown was created using BioRender. **(B)** Schematic workflow illustrating the quantification of cellular influences. Spatial objects were transformed and weighted using a spatial kernel, emphasizing cells that are in proximity. This approach captures variations in cellular influence quantification based on spatial organization despite the same cell counts within a region. **(C)** Downstream analyses included cellular neighborhood classification and evaluation of functional states.

Following cell type annotation, the FunCN model was applied. Similar to KNN-based or distance-based CN’s, we normalize the FunCN to represent the proportional influence exerted by each distinct cell type. However, unlike KNN or distance-based CN’s, FunCN does not create an arbitrary boundary for the cellular neighborhood and instead, considers the potential influence from all cells in the tissue, weighted by a rapidly decreasing spatial kernel. By implicitly incorporating both the proximity and local abundance of cell types, FunCN can reveal subtle differences in spatial organizations through distinct patterns of proportional influence—even when overall cellular composition is identical (**Figure 1B**). By associating the strongest influence from the nearest neighbors, this approach leads to a more precise representation of the cell-cell interactions.

The kernel function and width can be set to model either physical interactions between adjacent cells or diffusion of a secreted cytokine^24,25^ within a broader tissue region. The kernel bandwidth parameter determines the rate of decay of influence with distance. For instance, a smaller bandwidth places greater emphasis on the nearest cells and drastically decreases the influence as the distance increases. In contrast, a larger bandwidth results in a more gradual decrease in the influence over a broader radius, distributing weights more evenly across a larger area.

We then performed downstream analyses to identify groups of cells that shared similar local microenvironments. By leveraging spatial features derived from FunCN quantification, we clustered cells into distinct cellular neighborhoods (CNs), each representing a unique configuration of neighboring cell types and spatial proximity. Within each CN, we further examined cell state heterogeneity to understand how cellular behavior is shaped by spatial interactions within the TME (**Figure 1C**).

### Site-specific cellular compositions of primary and metastatic PDAC

To investigate differences in the cellular landscape between primary and metastatic PDAC, we examined site-specific cellular compositions. SOM clustering of protein expression profile was used to define cellular phenotypes. Among the resolved populations, we also observed an immune cluster that co-expressed markers of multiple immune lineages (CD4, CD8, DC-SIGN, CD86, CD206, and HLA-DR). Histological examination and raw images confirmed that this region contained a highly dense mixture of immune cell types, including dendritic cells (DCs, CD8^+^ T cells, and CD4^+^ T cells; **Figure S1B**). Further refinement of the clustering identified regulatory T cells, six distinct myeloid subtypes and seven stromal subtypes, each characterized by specific marker expression patterns (**Figure 2A, S1C-E**). When comparing the relative abundance of immune cells, both pancreatic primary tumors and liver metastatic lesions exhibited high proportions of CD68^+^ macrophages and CD15^+^ neutrophils. CD68^+^ CD163^lo^ CD206^lo^ (M_I) and HLA-DR^hi^ CD68^hi^ CD163^hi^ CD206^hi^ CD86^hi^ myeloid cells (M_II) were particularly high in abundance (**Figure S1F-G**). Out of all cell types, the collagen^+^ SMA/VIM^+/lo^ stromal subtype (Str_I, akin to myofibroblastic cancer-associated fibroblasts, myCAF) was the second most abundant cell type in both sites, following tumor cells. Comparison of cell type fractions between the two sites revealed that specific podoplanin^hi^ stromal subtypes (Str_VI and Str_VII) were more predominant in the pancreas, whereas Tregs, Str_II (collagen^+^ CD74^+^ SMA/VIM^+^ HLA-DR^+^, akin to antigen-presenting cancer-associated fibroblasts, apCAF), M_IV, and neutrophils were more enriched in the liver (**Figure 2B**). Additionally, M_I and Str_VI cell types were significantly enriched in the pancreas, whereas Str_II, M_IV, M_V, and tumor cells were more abundant in the liver (**Figure 2C**). These findings highlighted distinct immune and stromal compositions in primary pancreatic tumors versus metastatic tumors in the liver, warranting an in-depth evaluation of site-specific spatial interactions and functional implications. Cell type annotation was validated through the examination of cell masks colored by cell type at the pixel level (**Figure 2D**) and histological review of tissue cores.

**Figure 2.**
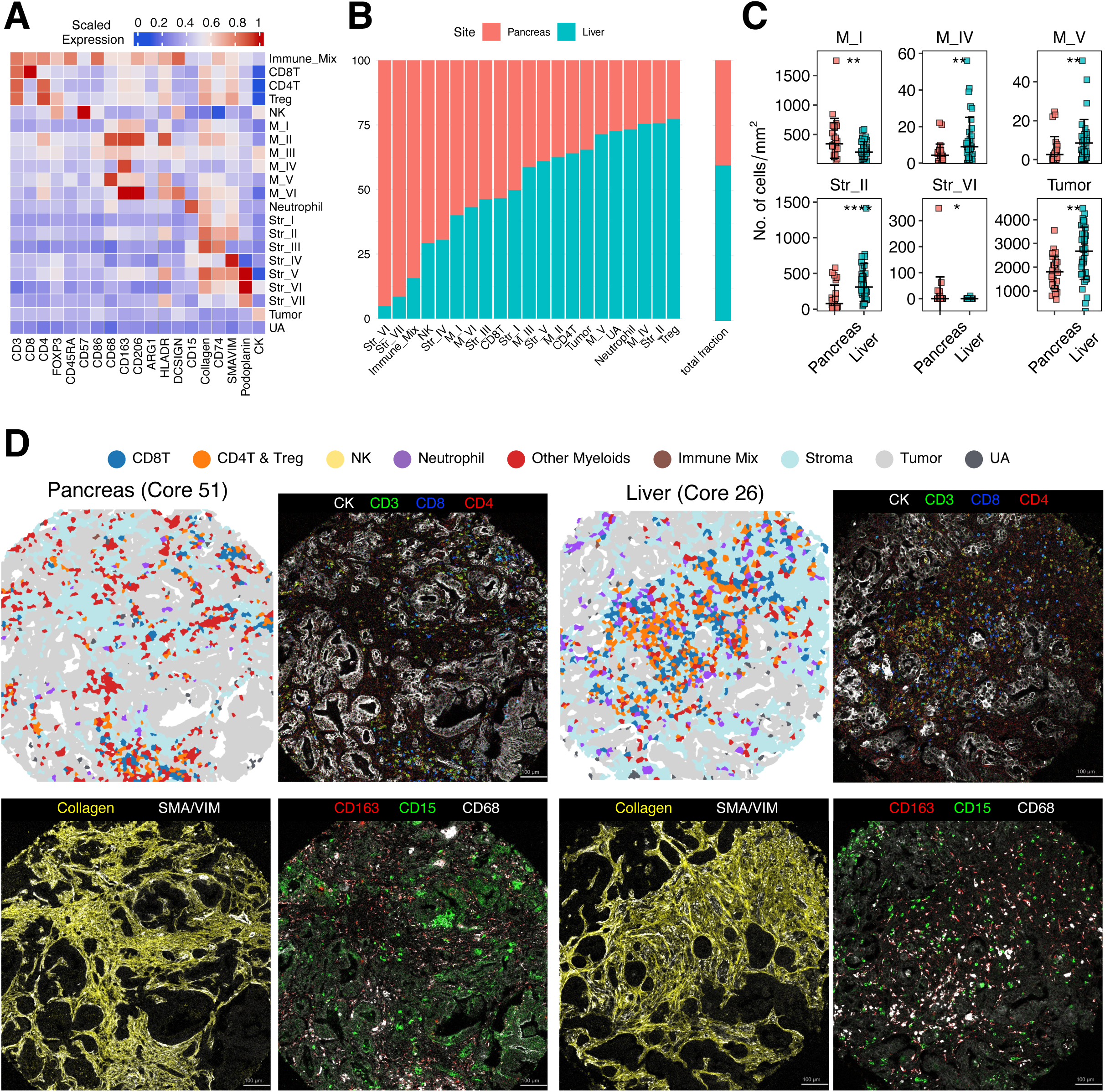
**(A)** Cell type phenotyping: Scaled expression profile of cell type signatures for each cluster is reflected as heatmap. **(B)** Proportional representation of cell types across sites and the relative abundance of each site shown as a fraction of the total cell population. **(C)** Cell types with significantly different densities (measured as the number of cells per mm^2^ area) between pancreas and liver. Differences were assessed using an unpaired Wilcoxon test. **(D)** Validation of cell type annotations. Representative cell mask images (left) from pancreatic and liver samples were compared to matched MCD images displaying relevant cell type markers. Scale bar: 100 µm.

### FunCN reveals enhanced resolution of spatial interactions in the tumor microenvironment

To validate our method, we compared the combined influence of CD8^+^ T and CD4^+^ T cells on tumor cells in core 51 (pancreas) with the spatial density plot of these two immune cell types in the same region. We observed high levels of combined CD8^+^ T and CD4^+^ T influence (shown in red, **Figure 3A**) on tumor cells located near areas where CD8^+^ T and CD4^+^ T cells were densely concentrated. We then explored how our FunCN approach detected distinct and subtle spatial patterns compared to the conventional k-nearest neighbor (KNN) method. Specifically, when quantifying cellular influence from the 10 nearest neighbors in core 51, FunCN method produced a wider, continuous range of values, whereas the KNN method provided only 0.1-step discrete increments (**Figure 3B**). The broader spectrum from the FunCN approach was especially pronounced at mid-range values, suggesting that in the regions of very low or very high cell frequencies, the frequency of cells was the main driver of cellular influence, whereas in the mid-range, distance-based weighting played a larger role.

**Figure 3.**
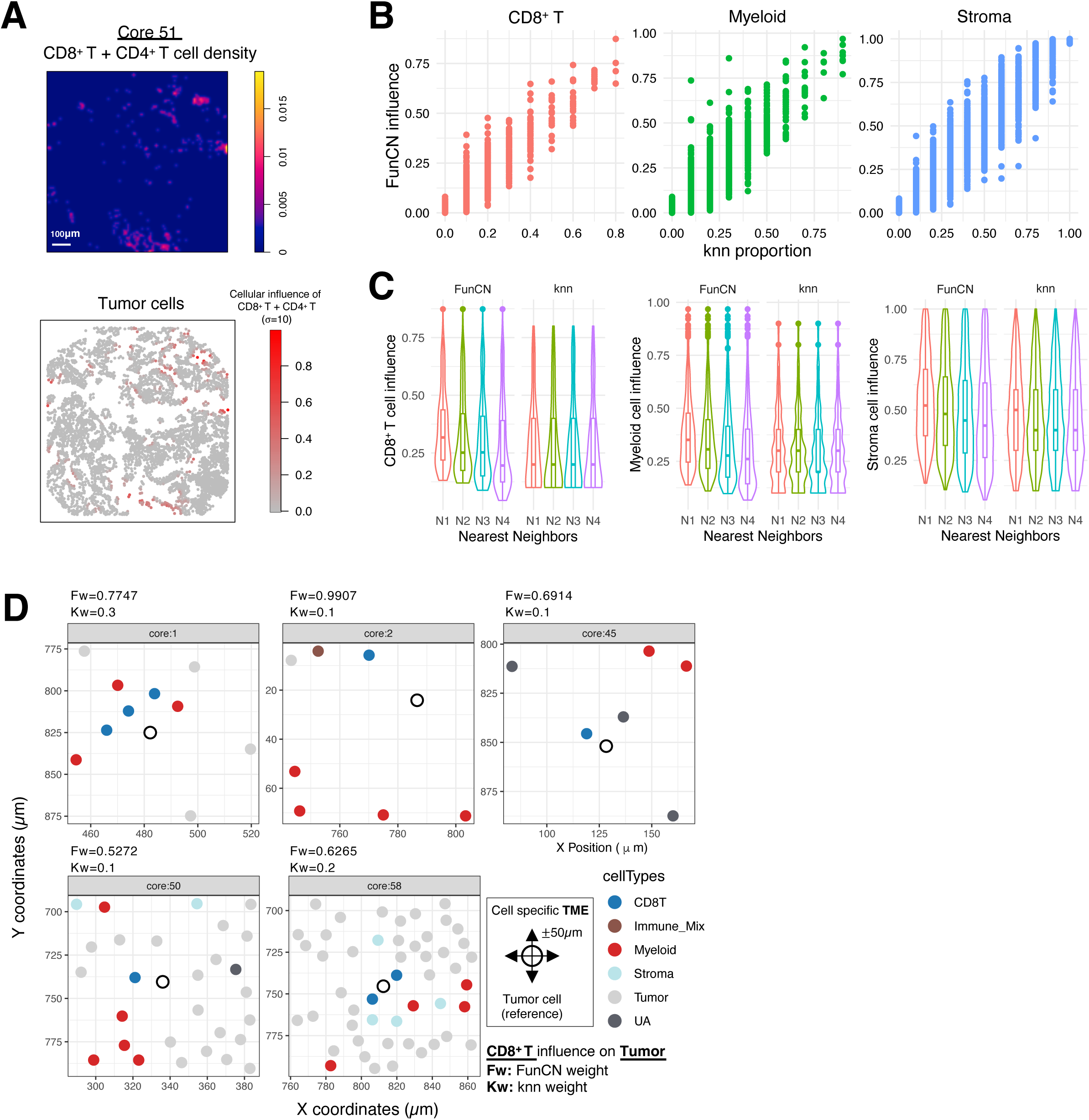
**(A)** Visualization of CD8^+^ T and CD4^+^ T cell influences on tumor cells within a representative core (Core 51). The top panel shows a spatial density plot representing T cell counts, while the bottom panel presents cellular influence values of tumors derived from the FunCN kernel weighting **(B)** Comparison of cellular influences from CD8^+^ T cells, myeloid cells, and stromal cells across all cells in the representative core, using the FunCN approach versus the KNN-based approach. For each discrete KNN spatial weight, continuous spatial weights were quantified using the FunCN. **(C)** Violin plots comparing FunCN and KNN methods for CD8T cells, myeloid cells, and stromal cells when these populations are among the N1, N2, N3, or N4 nearest neighbors. **(D)** Spatial organization of tumor cells showing the largest differences in spatial weights between the FunCN and KNN methods. Tumor microenvironment (TME) of a tumor cell within a ±50 μm radius are visualized, highlighting the increased emphasis on cellular influences when cells are in proximity.

Additionally, unlike the KNN approach that calculated cellular influence based on a fixed number of neighbors, our continuous weighting approach captured the effects of cells more precisely, even those that were beyond KNN’s cutoff. Our approach also ensured that rare but spatially proximal cells were not overshadowed by the higher overall frequency of other cell types in densely populated regions. Another key difference showed how weights were assigned based on neighbor order (n-th neighbors) (**Figure 3C**). Our FunCN approach showed a gradual decrease in CD8^+^ T, myeloid, and stromal cell influences as the neighbor order increased (with N1 being the closest and N4 being the furthest neighbor), whereas the KNN method, which did not consider distance, assigned equal weights across neighbors.

To examine the spatial organization of tissue specimens where the two methods would differ the most, we selected the top five tumor cells with the largest difference in CD8^+^ T cell influence between the FunCN and KNN methods. A ±50 µm window was set around each reference tumor cell. The results showed that CD8^+^ T cells in close proximity to the tumor cells were assigned higher influence values, with their influence considered more significant in our method (**Figure 3D**). This highlighted the ability of our method to capture interactions between tumor and immune cells.

Using our FunCN quantification method, we identified distinct spatial interactions in the primary pancreatic and metastatic liver TME (**Figure S2A-B**). Tumor cells at both sites showed limited cellular influence from other cell types, with only minimal contributions from M_I and Str_I. However, comparing the pancreas and liver, there was a greater influence of Str_II, Tregs, and neutrophils on tumor cells in the liver than in the pancreas, indicating distinct tumor-immune/stromal interactions in each of the sites. Notably, in the liver metastatic TME, stromal subtypes had more spatial influence from T cells. Neutrophil-CD8^+^ T influence was more enriched in the liver, whereas CD8^+^ T-other myeloid cell influence was more enriched in the pancreas. These site-specific enrichment of cellular influences, particularly those involving tumor cells and CD8^+^ T cells, suggested that the same cell types could have different functions depending on the tissue microenvironment and the neighboring cell types.

### Delineating distinct cellular neighborhoods around CD8^+^ T cells

We next conducted a more integrated analysis of the spatial influences to understand distinct cellular neighborhoods surrounding CD8^+^ T cells and tumor cells. To this end, CD8^+^ T cells were clustered based on their surrounding cell influences, yielding eight distinct cellular neighborhoods (CNs) via SOM clustering. These neighborhoods, derived from different patients and tissue sites, each showed unique cellular influences of multiple cell types, depicted by dot size indicating magnitude and color representing relative differences across CNs (**Figure 4A**). Two CNs, CN1 and CN2, were enriched for tumor cells. As expected, stromal cells were represented in CN2~6 but were mostly enriched in CN3 and CN6. CN5 was represented mostly by M_I myeloid cells but also had a stromal cell type, Str_I. Three CNs were enriched with T and myeloid cell subtypes at varying levels. CN4 was mixed with CD8^+^, CD4^+^, and regulatory T cells; CN7 was driven by primarily CD8^+^ T cells; and CN8 was mixed with CD8^+^ T and M_II type myeloid cells. Among these CNs surrounding CD8^+^ T cells, a large proportion of CD8^+^ T cells was found in proximity to neighborhoods enriched with CD8^+^ T cells (CN7), stromal (CN3, CN6), and M_I-type myeloid (CN5) cells. When comparing the two TME sites, we observed a predominance of CN7 over CN4 in the pancreas and the opposite for liver (**Figure 4B**), indicating greater influence from CD4^+^ and regulatory T cells on CD8^+^ T cells within the liver TME.

**Figure 4.**
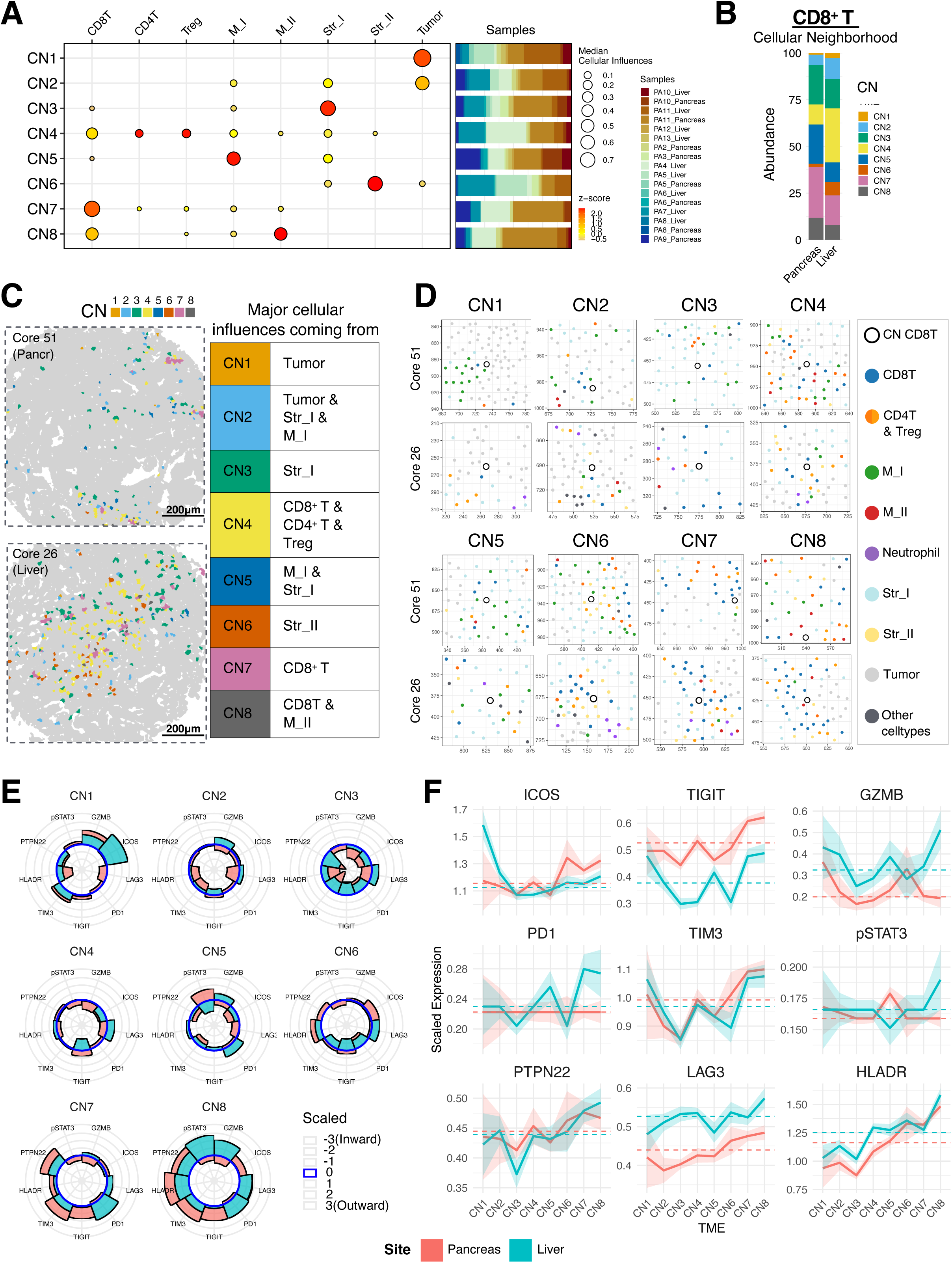
Cellular neighborhood (CN) analysis of CD8^+^ T cells. **(A)** CNs are depicted in a dot plot indicating the cellular influences comprising each neighborhood. Dot size represents the median cellular influence, while color indicates the z-score of differences across distinct CNs. A bar plot shows the sample fraction of each CN. **(B)** Stacked bar plot illustrating the abundance of CNs across different sites. **(C)** Representative images of CD8^+^ T cells in different CNs within a single core, demonstrating varying dominant cellular influences experienced by CD8^+^ T cells in that core. **(D)** For each CN, one CD8^+^ T cells was sampled from each representative core to validate its TME, highlighting relevant cellular influences as defined in the dot plot. **(E)** Analysis of CD8^+^ T cell functional marker expression, shown as radial plots indicating the relative z-scores of marker differences across CNs. **(F)** Line plot of median expression levels for functional markers with a shaded 95% confidence interval. The dashed line indicates the median expression of all CD8^+^ T cells in each site, with all CNs combined.

We further validated these CNs surrounding the CD8^+^ T cells by visual inspection. In each core, CD8^+^ T cells were assigned with different CNs based on their surrounding cell interactions (**Figure 4C**). To validate the spatial organization of CNs within CD8^+^ T cells, one cell per CN was sampled from each of two representative tissue cores 51 (Pancreas) and 26 (Liver), with X and Y coordinates zoomed within ±50 μm around the reference cells (**Figure 3D**). Across all CNs, CD8^+^ T cells exhibited a higher frequency and closer interactions with the specific cell types that characterize each neighborhood, consistent with the cellular compositions of the CNs.

We next explored the functional states of CD8^+^ T cells in relation to the CNs. Notably, CD8^+^ T cells in CN3, where Str_I cells exert the strongest influence and M_I myeloid cell type to lesser extent, exhibited lower expression of markers that denote activation (GZMB, ICOS, HLA-DR) in both the pancreas and liver (**Figure 4E**). This decreased expression of activation markers was also accompanied by lower expression of inhibitory markers (PTPN22, TIM3, and TIGIT). Taken together, our findings suggested that Str_I-type (collagen^+^ SMA/VIM^+/lo^ “myofibroblastic”) and M_I myeloid cells (CD68^+^ CD163^lo^ CD206^lo^ less activated macrophages) could diminish the activation of CD8^+^ T cells and impair their effector functions.

In contrast, CD8^+^ T cells in CN8, where M_II-type myeloid cells (CD68^hi^, HLA-DR^hi^, CD86^hi^, CD163^hi^, CD206^hi^ macrophages) exerted the strongest influence, exhibited higher expression of PTPN22, TIM3, TIGIT, LAG3, and HLA-DR compared to other CNs in both pancreas and liver, consistent with a state of immune exhaustion. However, CD8^+^ T cells in CN8 in the liver also expressed higher levels of GZMB compared to the pancreas, suggesting the presence of differential regulation by the tissue sites despite the similar cellular neighborhoods. Even though CD8^+^ T cells in the liver showed higher GZMB expression, we posit that other immunosuppressive signals in the microenvironment could inhibit overall T cell function^26^.

We also directly compared the functional marker expression level for each CN across both tissue sites. A line plot of median expression levels, with shaded areas representing 95% confidence intervals, illustrated trends in expression changes across CNs and highlighted differences between sites (**Figure 4F**). Interestingly, both sites exhibited remarkably similar trends for several functional markers, highlighting the critical importance of the surrounding cellular neighborhood in shaping the functional states of CD8^+^ T cells. For example, the expression trends for TIM3, HLA-DR, PTPN22, GZMB, and to a lesser degree LAG3 and TIGIT, were shifting together across the CNs regardless of the tissue site. Of note, CD8^+^ T cells in the liver, which were in close proximity to tumor cells with minimal stromal barriers, exhibited exceptionally high ICOS expression. Immune checkpoint marker expression also varied by site, with TIGIT levels elevated in the pancreas and LAG3 expression more prominent in the liver, indicating distinct immune exhaustion landscapes.

### CNs around tumor cells and their association with tumor cell expression of functional state markers

We applied the same approach to cluster tumors based on similar neighborhoods. Prior to CN classification, we first categorized tumors as either bulk or boundary tumor types, using a tumor cell influence cutoff of 0.7 (tumors with greater than and equal to 0.7 influence were classified as bulk, and those with less than 0.7, where 30% of the influence was attributed to other cell types, were classified as boundary tumors). We examined the boundary tumor cells because cellular influences from other cell types originate at the tumor boundary and propagate inward, allowing us to compare these influences and relate them meaningfully to the overall TME. This approach successfully distinguished boundary tumors, which exhibited greater interaction with other cell types, from bulk tumors, whose cellular influence primarily came from other tumor cells. (**Figure 5A-B**). Pancreas and liver sites had similar proportions of bulk and boundary tumor types (**Figure 5C**).

**Figure 5.**
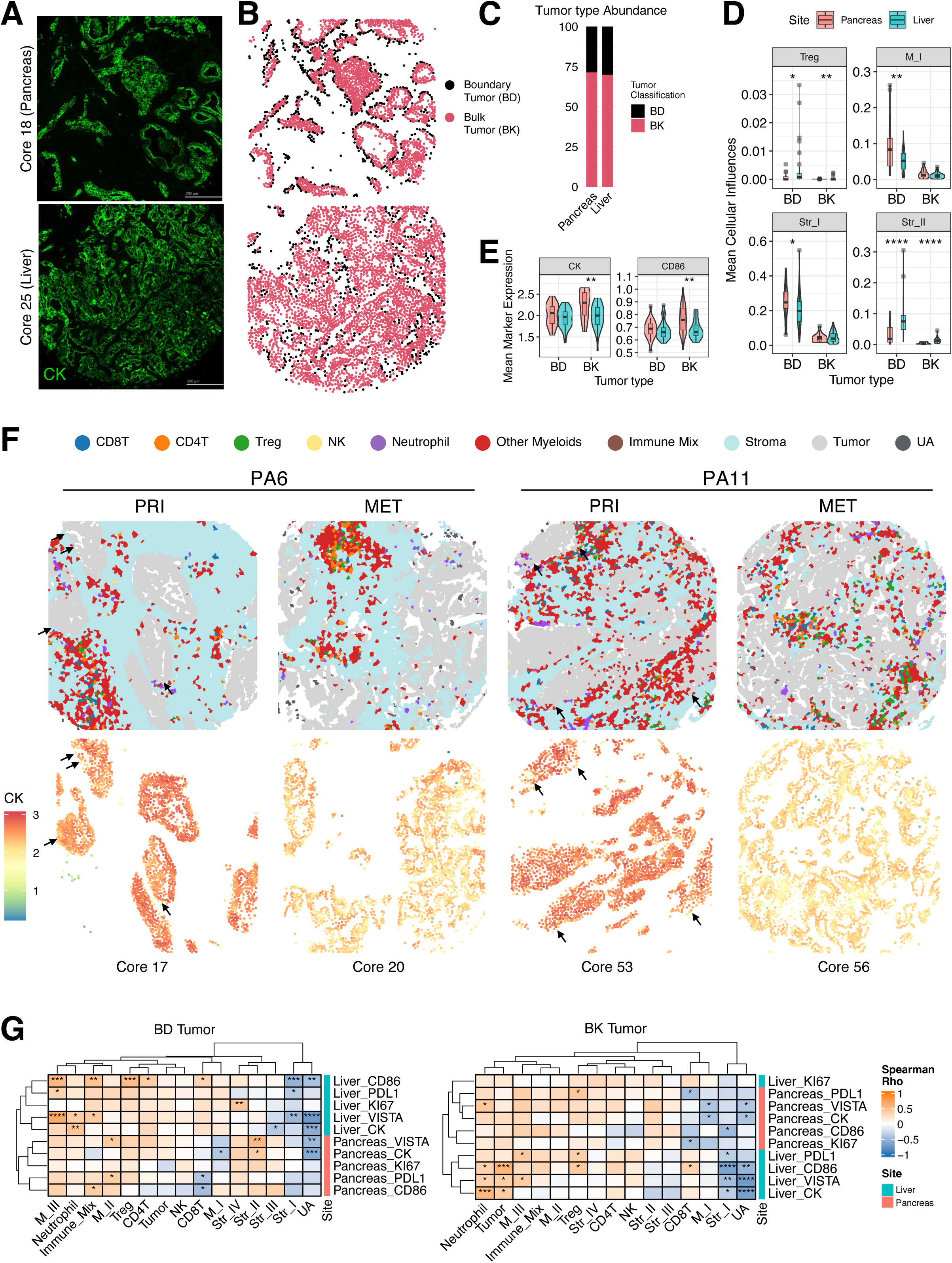
**(A)** Representative IMC images from the pancreas (core 18) and liver (core 25) showing the tumor marker CK. Scale bar: 200 µm. **(B)** Tumor cells were classified into boundary (BD) and bulk (BK) tumors based on a tumor influence cutoff of 0.7, with values below the cutoff classified as boundary tumors and values ≥0.7 classified as bulk tumors. **(C)** Bar plot illustrating the abundance of BD and BK tumor types across different sites. **(D-E)** Violin plots comparing the pancreas and liver for BD and BK tumor types. **(D)** shows significant differences in mean cellular influences, and **(E)** shows differences in mean expression of tumor markers CK and CD86. Statistical significance was assessed with an unpaired Wilcoxon test. **(F)** Cell mask images from matched primary and metastatic tumor sites (PA6, PA11) (top) and tumor cells colored according to CK expression levels (bottom). **(G)** Spearman correlation between mean cellular influence (mean per core) and mean tumor marker expression, with rho values shown as color gradients in the heatmap. Significant *p*-values are indicated (\**p* < 0.05, \**p* < 0.01, \*\**p* < 0.001, \*\*\**p* < 0.0001).

We identified 12 unique CNs in boundary tumors (**Figure S3A**), with major influences from neutrophils, other myeloid cells, stromal cells, and cells without sufficient marker expression for annotation (unassigned, “UA”, **Figure 1B**). Beyond tumor cells, stromal cells showed diverse patterns of influence across multiple CNs. Str_I (Collagen^+^ SMA/VIM^+/lo^, myCAF-like), in particular, exhibited varying magnitudes of influence in several CNs, with the strongest association observed in CN3. Meanwhile, other stromal subtypes were enriched in distinct CNs: CN7 (Str_II; Collagen^hi^ CD74^+^ HLA-DR^+^ SMA/VIM^+^, apCAF-like), CN4 (Str_III; Collagen^hi^ CD74^+^), and CN8 (Str IV; SMA/VIM^hi^). Proportion of tumor cells belonging to the Str_I-predominant CN3 was significantly higher in the pancreas, whereas the proportion in Str_II-predominant CN7 was significantly higher in the liver (**Figure S3B-C**). As for immune cells, M_I subtype myeloid cells (CD68^+^ CD163^lo^ CD206^lo^) exerted influence in CN9, CN10, and CN5, augmented by contributions from M_II (HLA- DR^hi^ CD68^hi^ CD163^hi^ CD206^hi^ CD86^hi^) in CN10 and M_III (DC-SIGN^+^, CD86^+^) in CN5 (**Figure S3A**). M_I-rich CN9 was significantly more represented in the pancreas than the liver (**Figure S3C**). While not statistically significant, neutrophil influence was particularly high in CN12 within several liver cores (**Figure S3C**). Consistent with these CN-based observations, tumor cells overall had stronger cellular influences from Tregs and Str_II in the liver, but more influences from M_I and Str_I cells in the pancreas (**Figure 5D**).

We next evaluated the expression of markers of functional states within the tumor cells across the CNs: CK for differentiation, PD-L1 and VISTA for immunosuppression, CD86 for costimulation, and Ki-67 for proliferation (**Figure S3D-E**). Tumor cells predominantly shaped by Str_I cells (CN3), and M_I cells (CN9) exhibited lower levels of these markers than those in other CNs (**Figure S3D**). Of note, tumor cells in the UA-predominant CN1 also expressed low levels of these markers. Based on histologic review, UA cells were mostly poorly differentiated tumor cells or other mesenchymal cells (**Figure S3F**). More specifically, in pancreas, CK expression was lower for CNs enriched with macrophages, CN9 and CN10 (**Figure S3D-E**), suggesting that macrophages promoted tumor de-differentiation and possibly epithelial-to-mesenchymal transition^27^. In addition, among the CNs, the neutrophil-enriched CN12 was associated with high tumor expression of immunosuppressive molecules PD-L1 and VISTA (**Figure S3D-E**), implicating the role of neutrophils in promoting tumor-mediated immunosuppression. Furthermore, when comparing tumor cells in general, bulk tumors in the liver showed lower expression of CK and CD86 compared to those in the pancreas (**Figure 5E**). CD86 expression was also lower in liver tumors, which plays active role in regulating the activity of T cells^28^.

Differences in CK levels across tumor cells from different sites and spatial organizations were also evident in the spatial visualization of tumors (**Figure 5F**). In the pancreas, tumors cells exhibited a more clustered formation, creating dense aggregations with overall higher CK expression. Metastatic tumor cells in the liver, on the other hand, demonstrated a more diffuse and less compartmentalized shape. Notably, in the pancreas, tumor cells with elevated CK were more abundant, particularly in clustered regions, whereas CK levels were relatively lower at boundary zones. Regarding immune interactions, both sites exhibited limited direct tumor– immune cell interactions, with most immune cells localized within the stromal compartment. However, in the liver’s diffused tumor structure, the stromal microenvironment appeared to recruit more Tregs and neutrophils, suggesting a more immunosuppressive TME that may contribute to immune evasion and reduced responsiveness to immune checkpoint therapies.

### Evaluating site-specific differences in the immune-tumor interactions and functional states

Building on these findings, we sought to further dissect site-specific immune regulation. To do this, we used Spearman correlation analysis to evaluate the pairwise relationships between cellular influences and functional states of tumor cells, using core-level averages of cellular influences and functional marker expression (**Figure 5G**). The spatial influences of two cell types, CD8^+^ T and M_I, showed opposing trends in relation to tumor functional marker levels in the pancreas and liver. In the pancreas, higher CD8^+^ T cell influence inversely correlated with tumor expression of immunoregulatory markers, resulting in lower CD86, PD-L1, and VISTA. In contrast, in the liver, greater CD8^+^ T cell influence correlated with increased expression of these markers. M_I exhibited similar relationships, though to a lesser extent. Other site-specific myeloid cell influences on tumor cells were also observed. Only in the liver, neutrophil influence correlated positively with elevated VISTA levels, while M_III cells (DC-SIGN^+^, CD86^+^) were significantly associated with increased expression of both VISTA and PD-L1. These varying immune-tumor relationships between the two sites suggested the presence of distinct immune regulatory mechanisms in each organ, influencing tumor immune evasion strategies differently. Namely, neutrophil/myeloid-driven enhancement of immunosuppressive environment was particularly notable in the liver.

To further investigate immunosuppression by VISTA, specifically as it related to neutrophil influence, we first classified the tumor cores into four inflammatory types based on the presence or absence of CD8^+^ T cell influence and PD-L1 expression, following the framework by Martin et al.^29^: Type I (“CD8 T+ PD-L1+”), Type II (“CD8 T-PD-L1-”), Type III (“CD8 T-PD-L1+”), and Type IV (“CD8 T+ PD-L1-”). These classifications provide a framework for TME archetypes, with Type I tumors harboring PD1/L1-inhibited T cell states, Type II representing immune ignorance, Type III exhibiting tumor-derived immune suppression, and Type IV suggesting the presence of alternative immunosuppressive pathways. Tumors were thus stratified into these four archetypes based on the median per-region values of CD8^+^ T cell influence and PD-L1 expression. We verified that tumor cells classified as “PD-L1+” exhibited higher PD-L1 expression (**Figure S4A**), and that “CD8 T+” classification exhibited a greater concentration of CD8^+^ T cells near the tumor cells (**Figures S4B**). Based on this classification, we observed that, specifically in the liver, neutrophil influence significantly correlated with VISTA expression in Type III TME (Spearman *R*=0.74, *p*=0.031) (**Figure S4B**). In the pancreas, although a significant correlation was observed in Type IV tumors (Spearman *R*=0.79, *p*=0.0098), the overall neutrophil influence was minimal (**Figure S4D**). This analysis supports the role of neutrophils in shaping an immunosuppressive environment in the liver tumors, particularly in Type III “CD8 T-PD-L1+” tumors, where immune evasion is mediated by tumor-intrinsic expression of immunosuppressive molecules. In line with these correlations, our analysis of the TCGA-PAAD data showed that patients in the top 20^th^ percentile for VISTA expression (VISTA-high group) had lower survival rates compared to those with lower VISTA expression (**Figure S4E**). Our finding highlights the impact of VISTA-mediated immunosuppression and implicates the potential need for VISTA-targeted therapies, especially in liver metastatic and neutrophil-enriched diseases, to improve patient outcomes.

Lastly, while more limited in sample size, we sought to account for patient-specific variability and evaluated the site-specific differences only using 6 primary-metastatic matched patient samples in a paired fashion. There was notable inter-patient variability, not only in immune cell influences on tumor cells but also in the expression of functional state markers by tumor cells (**Figure S4F-G**). We focused on patients exhibiting marked differences between primary tumors and metastases. For example, in patient PA11, the metastatic site showed a reduction in effector T cells (CD8^+^ T and CD4^+^ T cells) and NK cells, alongside an increased influence of Treg cells. This shift was accompanied by a reduction in CK levels and an increase in CD86 levels in the metastatic site. In PA6, a substantial reduction in tumor functional markers (CK, PD-L1, CD86, and VISTA) was observed in metastasis. The influences from other cell types (T cell, myeloid, and stromal) on tumor cells was low, indicating fewer interactions between the tumor cells and their microenvironment (stromal and immune cells). In PA10, as discussed earlier, a marked increase in VISTA-associated cell types, such as M_III (DC-SIGN^+^, CD86^+^) and neutrophils, was observed alongside elevated VISTA and PD-L1 levels in metastasis. Together, these findings highlight patient-specific immune adaptations, suggesting the need for personalized immunotherapeutic strategies.

## Discussion

In this paper we present FunCN, a spatial kernel-based model for quantifying cellular influence and defining cellular neighborhoods. FunCN incorporates both the frequency and the proximity of distinct cell types in the tissue to the target cell. Compared to prior art, FunCN applied to IMC data, established an enhanced characterization of the cellular organization and enabled a comprehensive mapping of the immune and stromal microenvironment of primary pancreatic tumors and liver metastases. FunCN is especially effective in accurately quantifying the strong localized influence of low-abundance cell types such as lymphoid and myeloid cells in the TME. It is important to emphasize that although FunCN quantifies the influence to represent the spatial organization of the TME, the functional state of cells remains inherently complex. For instance, determining whether a higher measured cellular influence translates to a more biologically meaningful impact on tumor behavior is still an open question. Future work could focus on integrating functional marker levels with proximity and abundance metrics to more comprehensively elucidate patient outcomes. Establishing the threshold at which cellular influences begin to significantly impact cell behavior is also a critical area for further investigation. Furthermore, different cell types can affect tumor function in distinct ways and their combined impact is challenging to fully disentangle. Our method to measure cellular influences could serve as an important foundation in developing predictive models of patient outcomes.

Applying FunCN based spatial analysis, we identified key immune and stromal interactions that may shape tumor immunogenicity. In pancreatic primary tumors, CD68^+^ myeloid cells were associated with reduced CK expression, suggesting a tumor-macrophage interaction that modulates epithelial differentiation. In contrast, in liver metastases, neutrophils and DC-SIGN^+^ CD86^+^ myeloid cells (M_III subtype) were linked to increased VISTA expression, reinforcing a myeloid-driven immunosuppressive microenvironment. The strong influence of neutrophils in mediating tumor immunogenicity through VISTA, particularly in CD8 T-PD-L1+ Type III tumors, suggests that immune checkpoint therapies relying on T cell activation may be less effective in these tumors. These findings suggest the need for therapeutic strategies targeting VISTA and neutrophil-mediated immune suppression to enhance immune infiltration and improve treatment responses in liver metastases.

Furthermore, LAG3 expression was consistently elevated in CD8^+^ T cells at the liver metastatic sites across most cellular neighborhoods. This finding aligns with a previous study^30^ reporting an increased proportion of LAG3^+^ CD8^+^ T cells in the liver. In contrast, at the pancreatic primary site, TIGIT expression was elevated in CD8^+^ T cells across the cellular neighborhoods. These results suggest the presence of tissue-specific drivers of distinct immunoregulatory axes. Similarly, our analysis uncovered opposing trends in CD8^+^ T cell influence on tumor functional states between pancreatic and liver tumors. In pancreatic tumors, CD8^+^ T cell influence was negatively correlated with functional markers (CD86, PD-L1, and VISTA), suggesting immune-mediated tumor suppression. However, in liver metastases, CD8^+^ T cell influence was positively correlated with these markers. Together, these findings indicate site-specific immune regulatory mechanisms, e.g., driven by molecular pathways and/or parenchymal signals, beyond what could be identified by the current antibody panel and IMC alone. These data provide new insights into why immune modulatory therapies that target different T cell suppressive pathways have different success rates depending on the organ sites hosting the tumors.

By integrating high-dimensional IMC spatial profiling with our FunCN model, our study provides a refined framework for understanding immune evasion mechanisms in pancreatic metastases. The identification of neutrophil-driven VISTA upregulation, macrophage-mediated CK suppression, and opposing CD8^+^ T cell influences across sites highlights the complexity of the metastatic immune landscape. These findings reinforce the need for site-specific, multi-target immunotherapeutic strategies, particularly those addressing VISTA, TIGIT, LAG3, and neutrophil-driven immunosuppression, to enhance immune activation and improve treatment outcomes in pancreatic cancer metastases.

## Resource availability

### Lead contact

Requests for further information and resources should be directed to and will be fulfilled by the lead contact, Won Jin Ho.

Address: 1650 Orleans St. CRB I Rm 488, Baltimore, MD 21287 Office: 410-502-5279, Fax: 410-614-9006

### Materials availability

This study did not generate new unique reagents.

### Data and code availability

Imaging mass cytometry data have been deposited at https://doi.org/10.5281/zenodo.15596960 and are publicly available as of the date of publication.

All original code has been deposited at GitHub (https://github.com/DeshpandeLab/Spatial_Influence) and is publicly available as of the date of publication.

Any additional information required to reanalyze the data reported in this paper is available from the lead contact upon request.

## Acknowledgements

This work was supported in part by the NIH/NCI T32CA193145 (J.W.L.), Break Through Cancer (L.D.W., A.D., W.J.H.); Maryland Cigarette Restitution Fund Research Grant to the Johns Hopkins Medical Institutions (FY25) (A.D.); The Lustgarten Foundation ‘A Translational Convergence Program of Personalized Immunotherapy for Pancreatic Cancer Patients at Johns Hopkins’ (E.M.J., A.D., W.J.H.), NIH S10OD034407 (W.J.H.), NIH/NCI P30CA006973 (W.J.H.), NIH/NCI U54CA268083 (L.D.W, W.J.H.), NIH/NCI P01CA247886 (E.M.J., W.J.H.).

## Author contributions

**Conceptualization**, Y.C., J.W.L., A.D., and W.J.H.; **methodology**, Y.C., J.W.L., S.M.S., A.G.H., X.Y., J.S., J.E.H., L.D.W., E.M.J., A.D., and W.J.H.; **software**, Y.C., J.W.L., S.M.S., A.D., and W.J.H.; **validation**, Y.C., J.W.L., A.D., and W.J.H.; **formal analysis**, Y.C., J.W.L., A.D., and W.J.H.; **investigation**, Y.C., J.W.L., S.M.S., A.G.H., X.Y., J.S., J.E.H., L.D.W., E.M.J., A.D., and W.J.H.; **resources,** Y.C., J.W.L., S.M.S., A.G.H., X.Y., J.S., J.E.H., L.D.W., E.M.J., A.D., and W.J.H.; **funding acquisition,** J.W.L., E.M.J., A.D., and W.J.H.; **data curation**, Y.C., J.W.L.,L.D.W., A.D., and W.J.H.; **writing – original draft**, Y.C., J.W.L., A.D., and W.J.H.; **writing – reviewing and editing**, all authors; **visualization**, Y.C., J.W.L., A.D., and W.J.H.; **supervision**, J.W.L., E.M.J., A.D., and W.J.H.; **project administration**, J.W.L., A.D., and W.J.H.

## Supplementary Figure legends

**Figure S1**

**(A)** Panel of antibodies used in the study. **(B)** H&E-stained image (left) of tissue cores with a high density of immune cell mix and corresponding MCD images (middle and right) colored by specific markers. Scale bar: 200 µm. **(C-E)** Scaled expression profiles of CD4T subtypes, other myeloid subtypes, and stromal cells subtypes **(F)** Stacked bar plot of abundances of each cell type across pancreatic and liver sites by patient (top) and by core (bottom). **(G)** Boxplot of proportion of each cell type (top) and the proportion of immune cells by site (bottom).

**Figure S2**

Analysis of cell-type-specific tumor microenvironment (TME). **(A)** Dot plot showing the mean cellular influences for each cell type across different sites, with dot size representing the mean cellular influence and color indicating scaled values relative to other cell types (scaled by column). Cellular influences below 0.01 and self-interactions were excluded. Differences in cellular influences between sites are highlighted with dashed boxes. **(B)** Radial plots illustrating differences in the TME of CD8T and tumor cells between the pancreas and liver. Bar height represents the log2 fold change (mean) between liver and pancreas (control), and color indicates the −log10(Bonferroni-adjusted) *p*-value from a Wilcoxon rank-sum test.

**Figure S3**

Cellular neighborhood (CN) analysis of Tumor cells. **(A)** CNs are depicted in a dot plot indicating the cellular influences comprising each neighborhood. Dot size represents the median cellular influence, while color indicates the z-score of differences across distinct CNs. A bar plot shows the sample fraction of each CN. **(B)** Stacked bar plot illustrating the abundance of CNs across different sites. **(C)** Violin plots comparing the proportion of tumor CNs in each core between sites. Statistical significance was assessed using an unpaired Wilcoxon test. **(D)** Analysis of tumor cell functional marker expression, shown as radial plots indicating the relative z-scores of marker differences across CNs. **(E)** Line plot of median expression levels for functional markers with a shaded 95% confidence interval. The dashed line indicates the median expression of all tumor cells in each site, with all CNs combined. **(F)** Representative H&E-stained image (Scale bar: 250 µm) (left) of tissue core with a high density of UA cells alongside the corresponding MCD image (right) showing CK (Scale bar: 200 µm) and cell mask (right) colored by cell types (UA cells shaded in dark gray).

**Figure S4**

**(A)** Box plot of average PDL1 expression level in tumor cells per core across different tumor classifications. Tumor types in each core are classified based on CD8^+^ T cell influence and PDL1 expression. **(B)** Representative MCD images for each tumor type, highlighting CD8 (CD8^+^ T cells) and CK (tumor) markers. Scale bar: 200 µm. **(C)** Box plot of average VISTA expression level in tumor cells per core across different tumor classifications. **(D)** Spearman correlation analysis of per-core average neutrophil influence and VISTA expression across different tumor types in pancreas (left) and liver (right). Each point represents a per-core average of tumor cells, with colored lines indicating the fitted regression for each tumor type. Shaded regions denote the confidence intervals. The correlation coefficient, R and p-value are displayed for significant associations in each site. **(E)** Kaplan-Meier survival curve comparing overall survival between patients with high (red, top 20^th^ percentile) and low (blue, bottom 20^th^ percentile) expression levels of VISTA (*VSIR*) gene in TCGA-PAAD dataset. The p-value from the log-rank test is shown. **(F-G)** Paired line plots of average values of immune and stromal cellular influences **(F)** and tumor-associated marker levels **(G)** in primary and metastatic tumors for each patient.

