## Supplementary figures and images for "Modeling cellular influence delineates functionally relevant cellular neighborhoods in primary and metastatic pancreatic ductal adenocarcinoma"

### Supplementary Figures

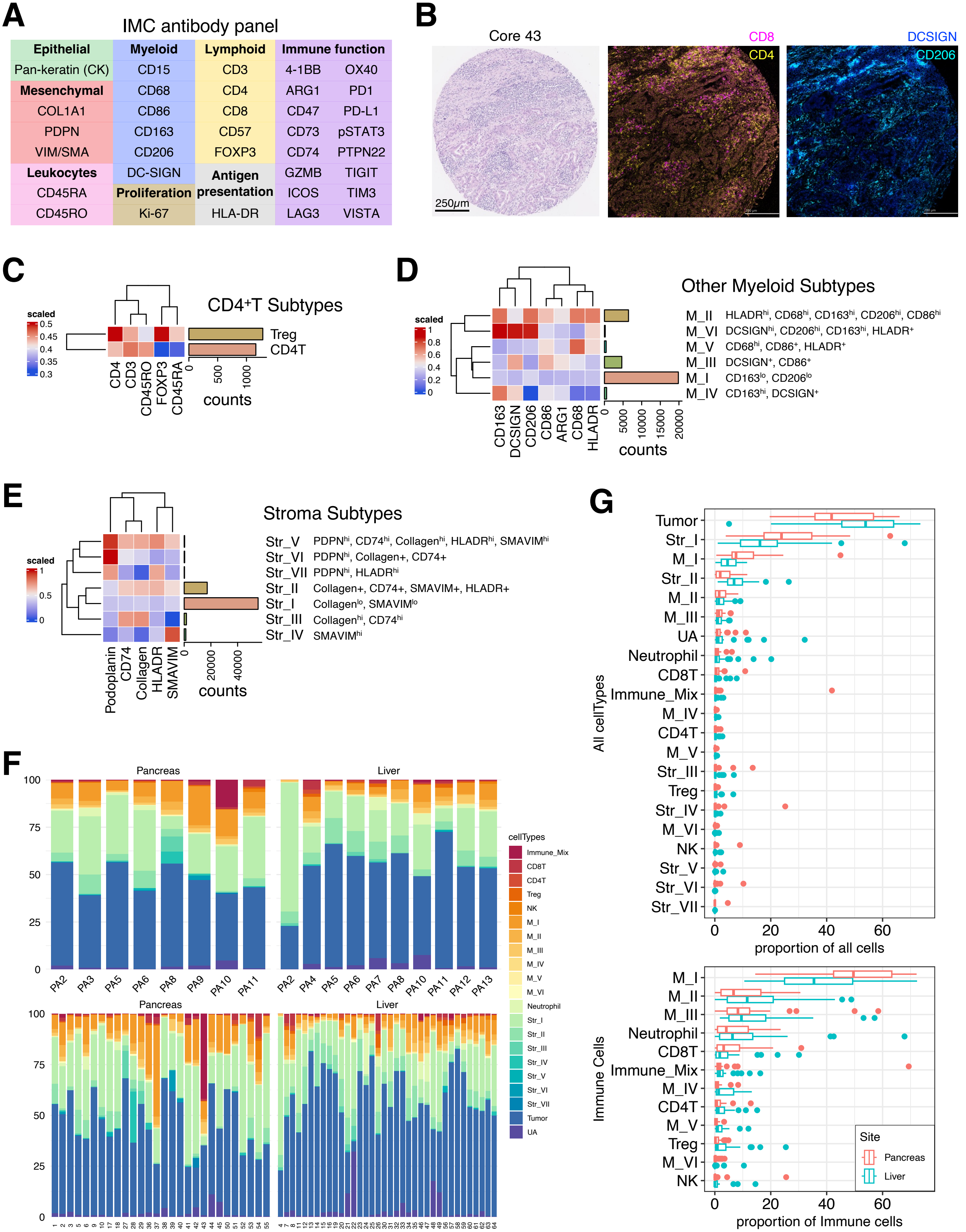


#### Supplementary Figure 1

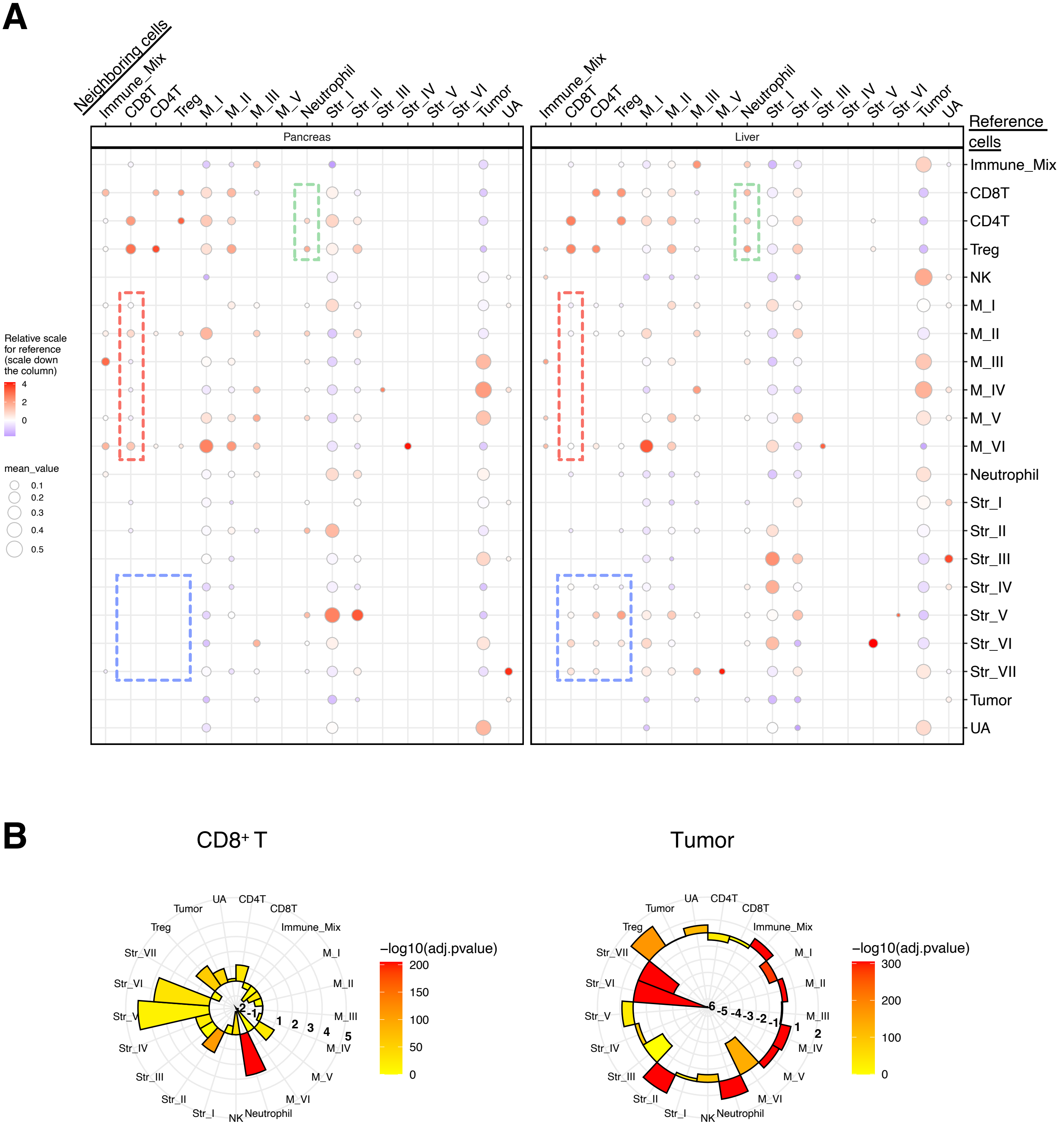


Supplementary Figure 2

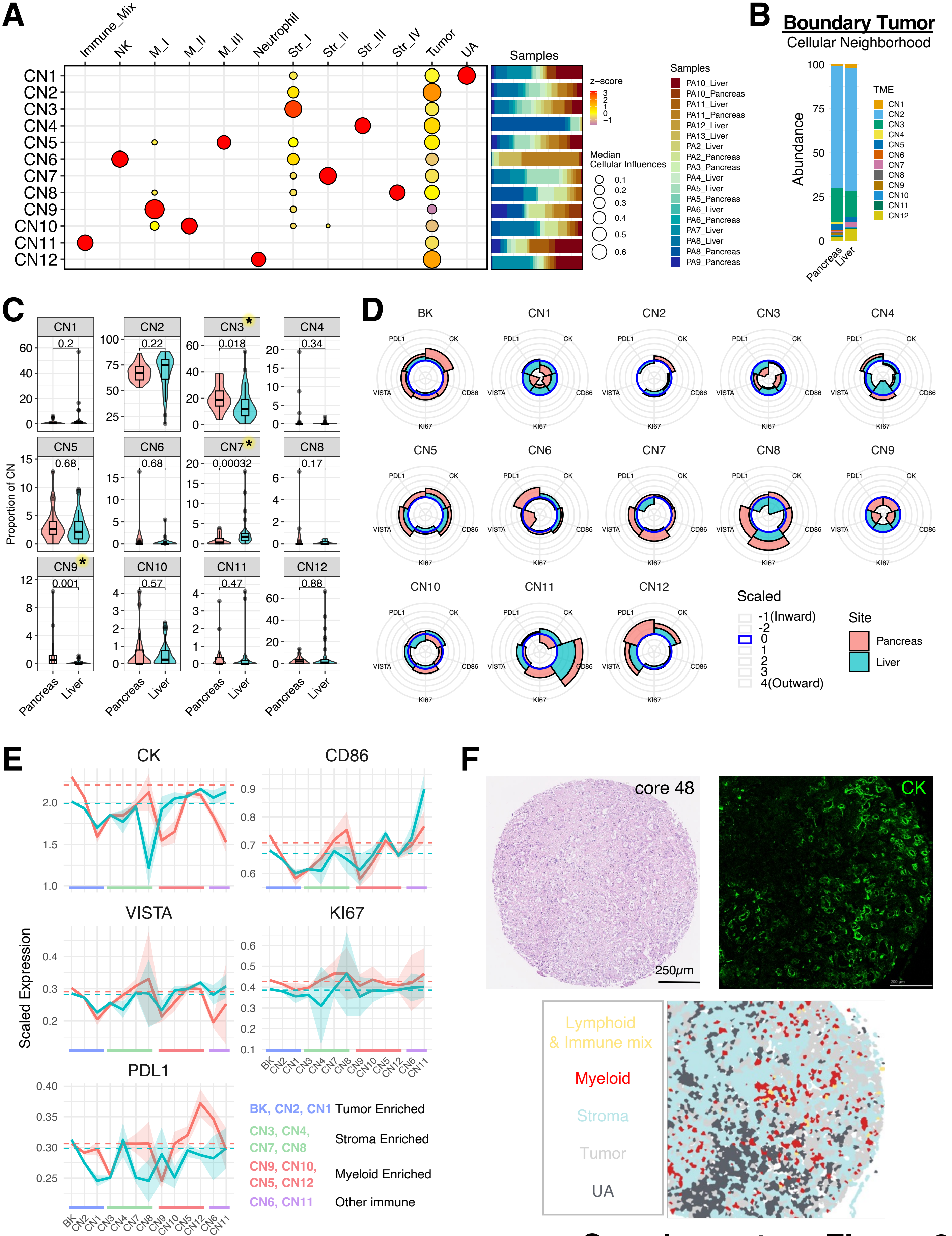


**Supplementary Figure 3**

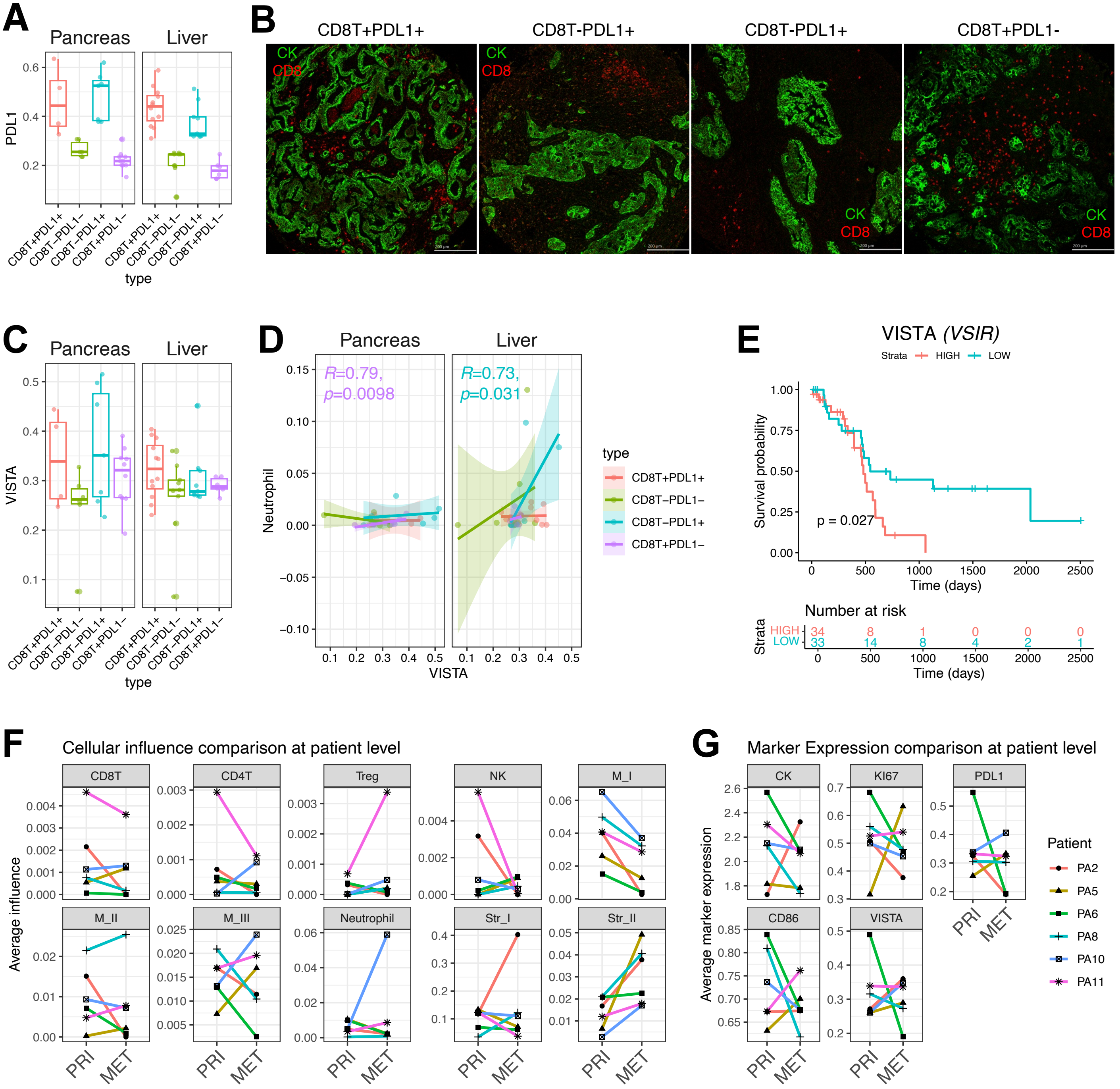


**Supplementary Figure 4**
